# Phenomenological model of transthyretin stabilization

**DOI:** 10.1101/2025.09.25.678455

**Authors:** Bartek Lisowski, Seweryn Ulaszek, Barbara Wiśniowska, Veronika Bernhauerová, Sebastian Polak

## Abstract

Transthyretin is a tetrameric transport protein whose monomers, when destabilized, can misfold and form amyloid fibrils, leading to serious diseases like transthyretin amyloidosis cardiomyopathy and neuropathy. While kinetic stabilisers such as tafamidis or acoramidis are designed to prevent tetramer dissociation, clinical data show a puzzling increase in TTR levels after treatment – an effect our study seeks to investigate by exploring possible underlying mechanisms.

Using a simple phenomenological model, we explore whether reduced dissociation alone accounts for this rise or if other mechanisms contribute. We propose that stabilisers may alter TTR clearance by slowing its cellular internalisation or degradation, or even by influencing its synthesis through pharmacological chaperoning. We also examine the role of monomer removal from circulation via re-association into tetramers or through other, possibly pathogenic processes. By integrating pharmacokinetic and pharmacodynamic data with experimental observations, our model provides fresh insights into TTR homeostasis and offers testable predictions for future research.

This study highlights the power of simplified, hypothesis-driven models in uncovering biological mechanisms – or, at the very least, in identifying key questions that remain to be answered.

## 1. Background

A central assumption in science is that insights from simplified laboratory systems should extend to real-world conditions. Natural observations inspire ideas, tested in controlled settings, and later re-examined in more complex systems. As Anderson noted, “more” is not necessarily better – only different [1]. Forgetting this leads to unnecessary experiments and overcomplicated models. Phenomenological models [2], [3]. Often overlooked in the life sciences – and rarely considered in medicine – these models can be built directly from existing data, without full mechanistic detail, and provide meaningful insight.

Our focus is transthyretin (TTR), a plasma protein mainly produced in the liver, also secreted by the choroid plexus and pancreas [4], [5], [6]. TTR transports thyroxine (T4) and, *via* retinol-binding protein (RBP), retinol. Functionally, it forms homo-tetramers that reversibly dissociate into monomers [7], [8], [9], [10]. In some variants – or even wild-type protein in the elderly – monomers misfold and aggregate into fibrils [11], [12] damaging tissues such as heart or nerves[13], [14], [15]. Mutant TTR causes hereditary amyloid cardiomyopathy (ATTRv-CM) or neuropathy (ATTRv-PN); wild-type aggregation leads to wild-type cardiomyopathy (ATTRwt-CM) [16].

Amyloid initiation remains unclear – misfolded monomers in blood, internalized tetramers, or intracellular events may be responsible. Nonetheless, stabilizers such as tafamidis and acoramidis slow disease by binding thyroxine sites, preventing dissociation and fibril formation [17], [18]. Subunit exchange assays, mixing labelled and unlabelled tetramers in plasma, quantify dissociation and reassociation kinetics, proving key to understanding stabilizer action.

More detailed models have incorporated drug competition with albumin [19], [20] and linked binding data, exchange assays, PK, and clinical results. All consistently show a >30% increase in circulating TTR after therapy [17], [21], [22]. A minimal model we proposed [23] suggested this rise cannot stem solely from slowed dissociation if monomers freely reassociate and are not degraded. By contrast, assuming slow reassociation and rapid monomer clearance readily explains the increase [20]. These opposing outcomes reflect uncertainties in monomer fate, highlighting the need for models that span plausible scenarios and yield testable predictions.

In this study, we examine how kinetic stabilisers [24] raise circulating TTR levels, explicitly considering reversible tetramer dissociation and monomer re-association as shown in subunit exchange experiments. We develop and parameterize PK/PD models of stabiliser action. Despite the biological complexity of TTR interactions with ligands such as retinol-binding protein, thyroxine, and albumin, we show that a phenomenological, data-driven approach can bypass many uncertainties. In particular, subunit exchange assays provide a direct link between stabiliser concentration and the effective reduction in tetramer dissociation, without requiring full knowledge of all binding partners. We focus on wild-type ATTR-CM as the reference case, while noting key differences in variant forms.

## 2. Methods

### 2.1 Transthyretin dynamics

We limit ourselves to the systemic circulation, in which TTR tetramers (*T*) are assumed to be constitutively produced in the liver and released to circulation at a constant rate *r*, as shown in Fig. 1. While in blood, TTR tetramers dissociate to and re-associate from monomers (*M*), with rates *k*_*d*_ and *k*_*a*_, respectively. It is also removed from blood (via internalization in various tissues, possible degradation and any other potential process leading to a decrease in transthyretin blood level) in a concentration-dependent manner with the rate *k*_*rem*_,_*T*_. Similarly, monomers are removed at a rate *k*_*rem,M*_. Under these assumptions, the dynamics of tetramers and monomers are governed by the following equations:

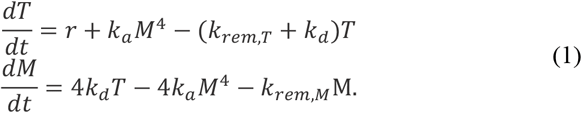

**Figure 1.**
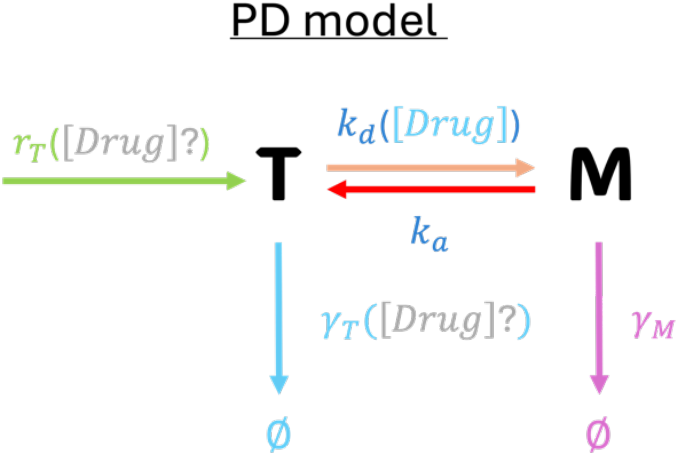
: Scheme of TTR minimal model, showing the transitions for tetramers, T, and monomers, M, together with corresponding rates (described in the main text). Possible in-direct effects of action of tetramer stabilisers are denoted as ([Drug]?).

Partial parametrization of this minimal model is possible due to excellent experiments, done *in vivo* by Jack H. Oppenheimer et al. back in 1960s [25] and more recently *in vitro* by Frank Schneider et al. [9] and R. Luke Wiseman et al. [26]. The former, namely Oppenheimer et al., in their Table 1, reports measured fractional elimination per day of total TTR tetramers (known previously as thyroxine-binding prealbumin, TBPA) for healthy and hospitalized human subjects, which can be used to estimate *γ*_*T*_. Knowing serum TTR steady state level and degradation, one can calculate tetramer production (and secretion) rate, *r*. The value reported in Table 1 was obtained under the assumption that *T*_*st*_ = *r*/*γ*_*T*_ to set a reference point, but this value should be treated with caution (see Sec. 3.1 for details). To convert from TTR mass to concentration, transthyretin molecular weight of 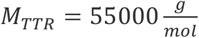 was used. We note that it is not always clear whether experimentalists and the kits they use measure only TTR tetramer concentrations or both tetramers and monomers together. Sekijima et al. [27] measured serum monomer concentration, which accounted for much less than 1% of total serum TTR. Therefore, unless otherwise stated, we interpret the experimental values of total TTR as equivalent to tetramer concentrations in the following discussion.

**Table 1.** Parameter values used in the minimal model of TTR dynamics (other parameters -*λ, α* and *γ*_*M*_ - are discussed in the main text)

| Parameter | Value | Reference |
| --- | --- | --- |
| $T_{st}$ | $6.44 \mu M$ | subject JO in Table 1 of [25] |
| $r$ | $0.1 \mu M \times h^{-1}$ | calculated as $r = \gamma_T T_{st}$ (but see Sec. 3.1) |
| $\gamma_T$ | $0.016 h^{-1}$ | subject JO in Table 1 of [25], parameter $K$ |
| $k_a$ | $360\,000 \mu M^{-3} \times h^{-1}$ | [26], estimated |
| $k_d$ | $0.0024 h^{-1}$ | data from Figure 5 in [28] for physiological temperature |

Tetramer dissociation rate has been measured in subunit exchange assays using ATTRwt patients’ plasma samples, also in physiological temperature [28]. This technique relies on an assumption, which is supported by observations, that monomer association into tetramers is relatively fast compared to dissociation. Wiseman et al. estimated *k*_*a*_ to be of the order of 10^20^*M*^−3^ × *s*^−1^ ≈ 360 000 *μM*^−3^ × *h*^−1^ [26]. All parameter values are shown in Table 1 and discussed in more details in the coming sections.

### 2.2 Pharmacokinetic Model

To parametrize PK model, we have used time-drug plasma concentration profile from [29] for the 7^th^ day of therapy with tafamidis single solid oral dosage formulation (61 mg free acid capsules), which was proven bioequivalent to the original tafamidis meglumine formulation (4 × 20 mg capsules once daily). The data were then fitted using a two-compartmental model, which was sufficient to mimic the pharmacokinetic profile of the drug (see SI):

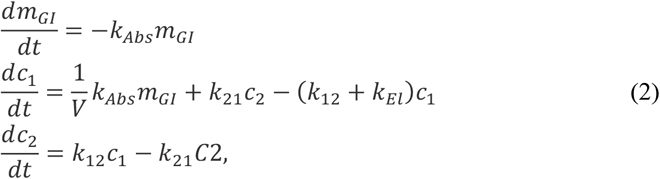

where *m*_*GI*_ is the mass of tafamidis in the gastro-intestinal tract, *c*_1_ and *c*_2_, are tafamidis concentrations in plasma and peripheral compartment, respectively. Transfer rates between compartments and elimination rate are explained in Fig. 2. Volume of the central compartment (i.e., blood plasma), needed for converting mass to concentration, is set to be equal to 3000 mL – 3/5 of the typical for human 5 L of blood [30]. Since Lockwood et al. report mass concentration, while stoichiometric relationships are defined in terms of molecular counts, in what follows we use molecular weight of tafamidis, 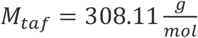, to convert from mass to molar concentrations.

**Figure 2.**
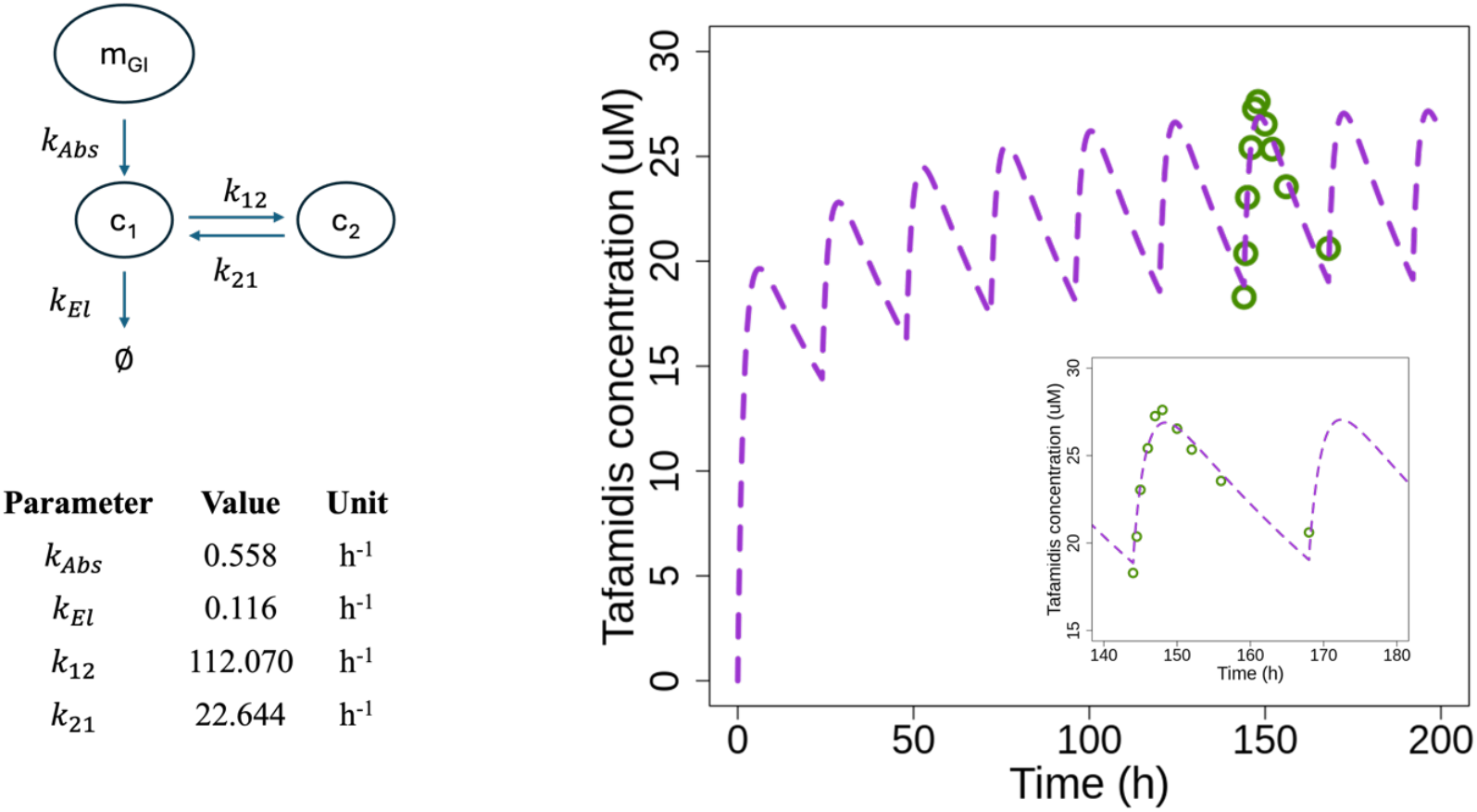
Pharmacokinetics of tafamidis is based on a two-compartmental model. Data used for parameter fitting are from [29].

All models were implemented in R Statistical Software (v4.4.2; R Core Team 2024). Nonlinear least-square function, nls(), and FME package were used to fit the parameters [31]. Plots digitization was done with the use of WebPlotDigitizer [32].

### 2.3 Subunit exchange and drug concentration-tetramer dissociation phenomenological relation

Phenomenological relation between TTR tetramer stability and tafamidis concentration in physiological temperature was established by Rappley et al. [28]. Using data from their Figure 5 we have fit the Arrhenius-like relation to capture the dependence of *k*_*d*_ on the plasma concentration of tafamidis, that is *c*_1_ in Eqs. (2):

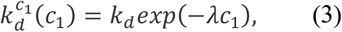

where *k*_*d*_ = 0.0024 *h*^−1^ is the rate of dissociation in the absence of drug (i.e., when *c*_1_ = 0), obtaining for the decay rate 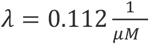, see Fig. 3. Such approach – fitting phenomenological relation given by Eq. (2) to data, without detailed knowledge between competing processes, off-target binding etc. – allows to describe tetramer dissociation with minimal set of parameters, among which only *λ* comes from fitting, while the rest – at least in principle – is well defined and can be obtained experimentally.

**Figure 3.**
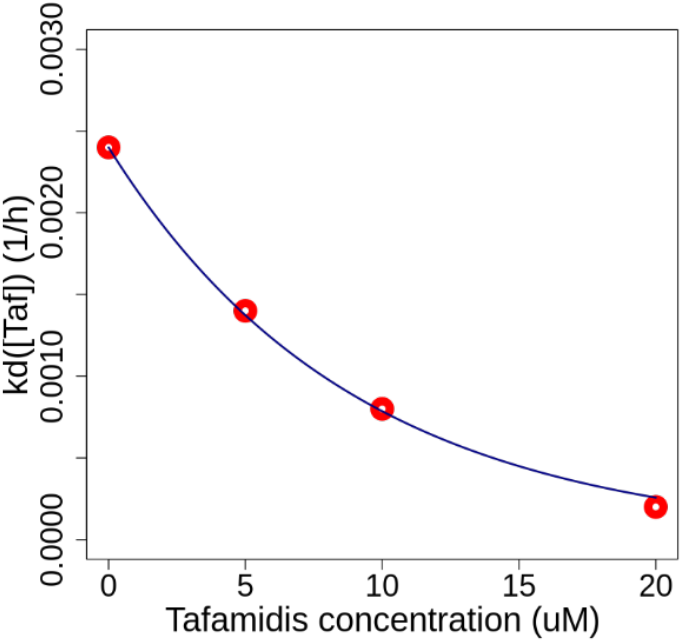
: Phenomenological relation between tetramer dissociation rate and tafamidis concentration. Red dots are experimental results from [28] and blue line is obtained by fitting Eq. (3) to the data, to find best-fit value of 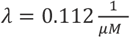.

## 3. Results

### 3.1 Simplicity of Pharmacodynamic Model Contrasts with Challenges in Parameter Estimation

At steady state, equations (1) reduce to:

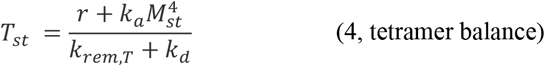

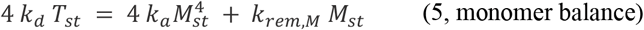

Here, *T*_*st*_ and *M*_*st*_ are steady-state tetramer and monomer concentrations, respectively.

Two processes remove monomers from the circulation: reassociation into tetramers 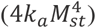 and elimination (*k*_*rem,M*_*M*). At steady state, the balance between monomer reassociation and degradation can be expressed by the dimensionless ratio

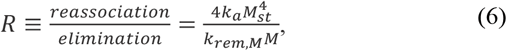

which quantifies their relative importance. When *R* is large (*R* ≫ 1), reassociation dominates and most monomers are recycled back into tetramers. In this case, the steady-state tetramer concentration approaches 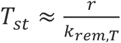 and is independent of *k*_*d*_. When *R* is small (*R* ≪ 1), degradation dominates, and most monomers are lost before they can reassemble. Here, 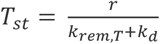, so reducing *k*_*d*_ immediately increases *T*_*st*_. In the intermediate regime (R∼1), both processes contribute comparably, and *T*_*st*_ shows only a partial and indirect dependence on *k*_*d*_ through its influence on the steady-state monomer level *M*_*st*_.

Having established the steady-state expressions for *M*_*st*_ and *T*_*st*_ in the reassociation-dominated, degradation-dominated, and intermediate regimes, we can now explore their practical consequences. The key point is that the analytical form of *M*_*st*_ and *T*_*st*_ depends on the relative magnitudes of the fluxes 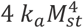 and *k*_*rem,M*_ *M*. Knowing several parameters from independent sources – for example, *T*_*st*_ and monomer-to-tetramer ratio *M*_*st*_/*T*_*st*_ ≪ 1% from human *in vivo* measurements, *k*_*rem,T*_ from tracer clearance data, *k*_*d*_ with and without stabilisers from subunit exchange experiments, and *k*_*a*_ from kinetic fits – allows us to insert these values into the regime-specific steady-state formulas and solve for the two remaining unknowns, namely the rates of tetramer synthesis, *r*, and monomer removal, *k*_*rem,M*_.

For the reassociation-dominated regime (*R* ≫ 1), the steady state for tetramers reduces to *T*_*st*_ ≈ *r*/*k*_*rem,T*_, implying *r* = *k*_*rem,T*_*T*_*st*_ and *k*_*rem,M*_ is negligible. In the degradation-dominated regime (*R* ≪ 1), the tetramer steady state is *T*_*st*_ = *r*/(*k*_*rem,T*_ + *k*_*d*_), so *r* = *T*_*st*_(*k*_*rem,T*_ + *k*_*d*_), and *k*_*rem,M*_ follows from the monomer balance. In the intermediate case (R∼1), one obtains, from the monomer balance, that 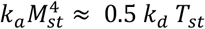, and, after inserting this to the expression for *T*_*st*_ and rearranging, *T*_*st*_ = *r*/(*k*_*rem,T*_ + 0.5*k*_*d*_), from which tetramer synthesis rate again can be recovered.

This analysis shows that, while steady-state formulas allow us to back-calculate *r*, the inferred value depends strongly on which regime is assumed. Direct experimental measurement of either the tetramer synthesis rate or the monomer removal rate *k*_*rem,M*_ would therefore be invaluable: by fixing one of these key parameters, and relying on those we already trust, we could distinguish between the reassociation-, degradation-, and intermediate-dominated regimes and thereby test or falsify some of the underlying assumptions.

### 3.2 Even perfect tetramer stabilisation accounts for only half of the clinical effect

Once the unknown parameters are determined for each regime, we can examine the hypothetical effect of perfect kinetic stabilization (i.e., when *k*_*d*_ → 0). In the degradation-dominated regime, the relative increase after cancelling out the possibility for tetramer dissociation is

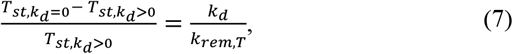

which, for the representative parameter set (*k*_*d*_ = 0.0024 *h*^−1^, *k*_*rem,T*_ = 0.016 *h*^−1^), corresponds to a maximum possible gain of about 15%. In the intermediate regime, the maximal gain is halved, while in the reassociation-dominated regime *T*_*st*_ is independent of *k*_*d*_, so no increase is possible. In each case, relative gain is independent of the inferred tetramer synthesis rate *r*. These theoretical bounds make it clear that achieving a 30% increase in total TTR purely through dissociation suppression would require parameter values outside those typically observed, or additional mechanisms beyond simple kinetic stabilization.

### 3.3 Pharmacokinetic Model Accurately Predicts Tafamidis Plasma Concentration Profile

Figure 2 shows tafamidis concentration profile and parameter values predicted by the PK Model described in Sec. 2.2. Two compartmental model, which distinguishes between concentrations in the central *c*_1_ (i.e., blood plasma) and peripheral (i.e., tissues or rest of the body) compartments corresponds well with *in vivo* data from [29]. Above all, the predicted steady state concentration of tafamidis has a value of around 25 *μM*, what stays in agreement with drug’s specification [33].

Further presentation of the results requires a comment. Here, we limit the use of the PK model to reproducing the total concentration-time profile in plasma, without trying to explain the estimated parameter values. This approach is justified for at least two reasons.

First, TTR is not the only plasma protein capable of binding tafamidis or other stabilisers. For example, while albumin has an order of magnitude lower affinity for tafamidis than TTR, it is at least ten times more abundant. As a result, competition between targets ensures that not all plasma tafamidis is available to bind and stabilize TTR [19].

Second, not all TTR in plasma is fully accessible for stabiliser binding due to its role as a transport protein. Each stabiliser’s molecule can bind only to one of the two thyroxine (T4) binding sites on a TTR tetramer. Although most plasma T4 is transported by thyroxine-binding globulin (TBG), up to 15% of TTR molecules have T4 bound to these sites, making them unavailable for stabilisers [34]. Similarly, while the RBP-retinol complex does not sterically block other ligands from binding to the T4 sites, its binding kinetics likely differ from those of an unoccupied TTR tetramer [35], [36], [37], [38].

Incorporating these and other molecular details into a physiology-based pharmacokinetic (PBPK) model would surely enhance our understanding of the system. But parametrizing such a model is inherently challenging due to limited available data. This is where a phenomenological approach, paired with the *right* data, provides an effective means to make meaningful progress.

### 3.4 Subunit exchange assay eliminates the impact of known and unknown unknowns

The subunit exchange assay was developed to evaluate the effects of kinetic stabilisers in various types of samples, with its primary advantage being the ability to perform reliable measurements in blood plasma from both healthy individuals and patients with TTR amyloidosis. By tracking the rate at which monomers exchange between labelled and unlabelled tetramers, as well as the distribution of tetramers with different combinations of labelled and unlabelled monomers (e.g., 0 labelled – 4 unlabelled, 1 labelled – 3 unlabelled, etc.), as a function of a known stabiliser concentration added to donor plasma, it is possible to establish a phenomenological relationship between the tetramer dissociation rate (*k*_*d*_) and the stabiliser concentration, as shown in Fig. 3.

The strength of this approach lies precisely in its phenomenological nature: for a known concentration of a drug added to blood plasma – with all its known or potential targets – one measures the dissociation rate of the TTR tetramer. How much of it is bound to tafamidis, T4, or albumin instead of TTR does not matter. This is because the result is exactly what is needed to integrate pharmacokinetics (which relies on measuring only the total drug concentration in plasma, without distinguishing whether it is bound to TTR, other proteins, or unbound) with pharmacodynamics (which focuses on the impact of tetramer-stabiliser binding).

We use tafamidis as an example of a stabiliser due to the availability of data from subunit exchange experiments conducted at physiological temperature [28]. However, this approach is generally applicable to all known and clinically significant stabilisers [19] (see also [39] and [40] for further discussion of the method).

Thus, for an individual patient, performing subunit exchange experiments across several tafamidis concentrations allows fitting of Eq. (3) and direct estimation of the effective decay rate *λ*. Combined with the patient’s pharmacokinetic profile described even by a simple two-compartment model, this enables prediction of tafamidis concentrations over time and, consequently, the extent of TTR stabilization. Because the assay already incorporates the net effects of off-target binding, ligand competition, and variable drug occupancy, these predictions can be made without detailed knowledge of underlying molecular interactions, providing a practical basis for individualized therapy planning.

## 4. Discussion and conclusions

In this paper, we stressed the value of drawing maximum insight from existing data, supported by simple quantitative reasoning, while also pointing to key unresolved questions. One central issue is the role of TTR monomers in blood. Their concentration, *in vitro* and *in vivo*, represents only a small fraction of total TTR, dominated by tetramers. Yet this observation alone reveals little about the underlying dynamics, much like early misconceptions about HIV replication, which were overturned when Ho, Perelson, Bonhoeffer, Nowak and others showed that viral steady states arise from a balance of production and clearance, leading to the optimization of existing therapies and the development of new ones [41], [42].

An analogous approach is needed for TTR. Monomer levels reflect tetramer dissociation opposed by reassociation and elimination, the latter including degradation, amyloid deposition in vessels, or translocation into tissues, collectively described by *γ*_*M*_*M*. It is generally assumed that elimination dominates, since reassociation depends on the fourth power of monomer concentration, making it negligible compared to the linear elimination term. This not only determines tetramer steady state but also raises crucial questions about the fate of circulating monomers.

The key uncertainty is that monomer clearance from blood has never been measured directly. By analogy with proteins of similar size, rapid removal seems likely, but this remains unproven. Recent work suggests that dissociation and misfolding of TTR monomers may be constrained by molecular crowding[43]. Protein concentration in healthy human plasma is around 80 g/L. Using the (phenomenological) relation between molar concentration and mean molecular separation in nanometres, *d* = 1.18 *C*^−1/3^, we estimate that the average distance between plasma proteins is ∼15 nm [44]. This is only about three to four times the diameter of a TTR monomer (4–5 nm), making it unlikely that dissociated monomers can diffuse far without rapidly encountering other proteins – or each other – suggesting reassociation may be favoured. At the same time, recent reports indicate that amyloid deposits can already form within blood vessels [45]. If the affinity of monomers for such deposits were very high, and the likelihood of encountering them substantial, the “stabilizing” influence of molecular crowding would of course be diminished.

These considerations highlight the importance of analysing the time scales that govern monomer fate after tetramer dissociation. Future studies should measure monomer abundance, tissue penetration, and clearance in health and disease. Notably, other amyloid-forming proteins also exist in oligomeric states [46], [47], [48], [49]. Whether such non-linear concentration dependence is a general hallmark of amyloidoses or a coincidence remains an open question.

TTR concentration in blood is one of the few quantitative endpoints used to assess pharmacological stabilisation, though its role as biomarker or prognostic remains debated [50], [51], [52], [53], [54]. In ATTRwt and ATTRv patients, TTR concentration has been observed to increase by more than 30% after treatment initiation [17], [21], [22]. Pharmacodynamic models must account for this effect, yet reduction of tetramer dissociation alone may not suffice in an open system.

*In vitro*, lowering dissociation stabilises tetramers under mass conservation [23]. *In vivo*, however, TTR is continually synthesised and cleared by degradation or tissue uptake. Variant TTR removal after liver transplant follows exponential decay, consistent with model predictions [55].

Internalisation adds another layer. Sousa and Saraiva [56] showed that uptake is cell-specific and modulated by ligands: T4 accelerates it, while RBP slows it. Because ligand or drug binding alters TTR conformation, receptors likely sense these changes [57], [58]. Although untested, it is plausible that stabilisers such as tafamidis or acoramidis also affect internalisation—and possibly other rates as well [59].

Sousa and Saraiva also showed that TTR variants differ markedly in internalisation: the hyper-stable T119M was taken up faster than the amyloidogenic V30M, wild-type TTR, or the highly amyloidogenic L55P, which was barely internalised at all [56]. This highlights how conformational changes strongly affect clearance. Our earlier model assumed stabilisers act only by reducing tetramer dissociation, but it seems equally plausible that binding also alters elimination rate.

Stabilisers target the same tetramer sites as thyroxine, which itself stabilises TTR, and were inspired by the hyper-stable T119M [60], [61]. Notably, both T4-bound TTR and T119M are internalised faster than TTRwt [56]. Whether tafamidis or acoramidis have similar effects remains unknown, but such measurements are crucial to understand their impact on TTR turnover. By analogy, stabilisers might even accelerate clearance rather than reduce it.

A further question is whether internalisation always leads to degradation. Sousa and Saraiva observed that hepatic degradation progressed linearly over hours, while internalisation quickly saturated [56]. Thus, uptake and degradation are at least partly decoupled, and reduced *γ*_*T*_ may result once internalisation plateaus. Another possibility is that stabilised tetramers are exocytosed back into circulation, effectively lowering elimination. Though unproven, some evidence suggests cells may preferentially secrete stable tetramers rather than degrade them.

Alternatively, could stabilised tetramers be exocytosed back into the circulation, thereby reducing net elimination rate, *γ*_*T*_? Though untested, evidence suggests cells can favour release of stable tetramers during synthesis and secretion. It is therefore crucial to study tetramer uptake, endothelial transit, degradation, and tissue accumulation, and to determine how stabilisers modulate these processes. Transcytosis could be tested by saturating cells with TTR, then measuring its release into fresh medium. Equally important is understanding TTR production and whether feedback mechanisms exist. Pharmacological chaperoning has been reported [59], but it does not explain how circulating TTR might regulate its own hepatic synthesis – unless internalised TTR acts as an environmental signal, akin to lactose in the lac operon. If so, the assumption of constitutive production would need revision.

Transthyretin amyloidoses pose a major medical and economic burden. Even with improving therapies, late diagnosis and aging populations will sustain the challenge. Advances in models and treatments have clarified ATTR, yet fundamental physiology – TTR secretion, clearance, internalisation, and tissue penetration – remains poorly understood. Closing these gaps is vital for future progress.

## Abbreviations

ATTR: transthyretin amyloidosis
ATTRv-CM: hereditary amyloid cardiomyopathy
ATTRv-PN: hereditary amyloid neuropathy
ATTRwt-CM: wild-type cardiomyopathy
PBPK: physiologically-based pharmacokinetics
PD: pharmacodynamics
PK: pharmacokinetics
RBP: retinol binding protein
T4: thyroxi
TBPA: thyroxine-binding prealbumin
TTR: transthyretin

## 5. Acknowledgements

This research was co-funded by the European Union within the VITAL Horizon Europe project (Grant nr 101136728). BL and VB are supported by a grant from the Priority Research Area qLIFE under the Strategic Programme Excellence Initiative at Jagiellonian University under the agreement 06/IDUB/2019/94. VB acknowledges the support of UNCE, project number UNCE/24/MED/008 and the project New Technologies for Translational Research in Pharmaceutical Sciences /NETPHARM, project ID CZ.02.01.01/00/22_008/0004607, co-funded by the European Union.

## 6. Supplementary Information

SI in the form of Colab Notebook are available under the link: https://colab.research.google.com/drive/1l1f2wVVm_tFYK8UNkCecHPgAt3Fd-UCW?usp=sharing

